# Evidence for Early Evolution of Sulfated Peptide Signaling in Plant Development

**DOI:** 10.1101/2025.11.07.687175

**Authors:** Devin V. Tulio, Alexandra M. Shigenaga, Shu-Zon Wu, Pamela C. Ronald, Magdalena Bezanilla

## Abstract

In plants, the cell wall fixes the position of each cell; therefore, during development, plants rely on cellular proliferation and expansion for tissue patterning and organ formation. How plant cells communicate with neighboring cells to coordinate expansion for properly patterned tissues and organs is not well understood. In seed plants, organ growth is known to be modulated by sulfotyrosyl peptide signaling. Here, we report that the activity of TYROSYL PROTEIN SULFOTRANSFERASE (TPST), which is responsible for the post-translational modification of sulfotyrosyl peptides, is essential for expansion during development in the non-vascular plant, *Physcomitrium patens*. Plants that harbor a null mutation in the gene encoding TPST (Δ*tpst*) were smaller, formed fewer caulonemal filaments, and were unable to form expanded gametophores. In Δ*tpst* multiple aspects of gametophore development were affected, including the first division of the gametophore initial, as well as reduced rates of cell division and expansion. Mutational analysis of *P. patens* TPST identified the residue Histidine 124, a candidate catalytic residue, as essential for TPST function. Notably, addition of the sulfated signaling peptide, PSY1 from either *P. patens* or *Arabidopsis thaliana*, rescued all Δ*tpst* developmental deficits. Taken together, these data suggest that TPST functions to sulfate PSY, and this activity is necessary for plant growth and development. Furthermore, since addition of AtPSY fully rescues Δ*tpst* and PpPSY promotes root elongation in Arabidopsis and rice, these findings suggest that PSY signaling is evolutionarily conserved.

## Introduction

In multicellular organisms, development involves cell proliferation, expansion, and differentiation. Unlike animal cells, plant cells are unable to undergo cell migration due to their encasement in a semi-rigid cell wall. Thus, the ultimate body shape depends on patterns of cell division and cell expansion (Cosgrove, 2005). How plant cells coordinate expansion with neighboring cells to shape tissue development and organ formation is not known.

Tyrosine-sulfated peptide signaling regulates plant growth, development, and immune response (Kaufmann and Sauter, 2019). A member of the <u>P</u>LANT PEPTIDE-CONTAINING <u>S</u>ULFATED T<u>Y</u>ROSINE (PSY) family, mature PSY1 is an 18-amino acid tyrosine-sulfated peptide (Amano *et al*., 2007). Exogenous PSY1 at nanomolar concentrations enhances cellular expansion and proliferation in Arabidopsis (Amano *et al*., 2007) and overexpression or exogenous application of PSY1 promotes root elongation in Arabidopsis and rice (Amano *et al*., 2007; Pruitt *et al*., 2017; Ercoli *et al*., 2024; Shi *et al*., 2024). Other characterized sulfopeptide hormones include <u>P</u>HYTO<u>S</u>ULFO<u>K</u>INE (PSK), a 5-amino acid tyrosine-sulfated peptide that functions with the plant hormones auxin and cytokinin to mediate cell cycle reentry (Matsubayashi and Sakagami, 1996; Matsubayashi *et al*., 1999), <u>R</u>OOT MERISTEM <u>G</u>ROWTH <u>F</u>ACTORS (RGFs), and <u>C</u>ASPARIAN STRIP INTEGRITY <u>F</u>ACTORS (CIFs) (Matsuzaki *et al*., 2010; Doblas *et al*., 2017; Nakayama *et al*., 2017). However, the molecular mechanisms that link tyrosine-sulfated peptides to cell expansion are not well understood.

In plants, sulfated signaling peptides are processed and post-translationally modified from precursor peptides. The prepropeptide amino-terminal signal peptide is cleaved during secretion in the endoplasmic reticulum, and then it moves to the Golgi for tyrosine sulfation and other modifications as appropriate. Propeptides are then transported, likely to the apoplast, and become mature sulfated signaling peptides when they are cleaved at the amino- and carboxyl-termini by largely unknown proteases (Komori *et al*., 2009; Kaufmann and Sauter, 2019).

Sulfotyrosine (sTyr) residues are formed by TYROSYL PROTEIN SULFOTRANSFERASE (TPST) in the Golgi and are critical for certain protein-protein interactions in animals and plants (Stewart and Ronald, 2022). TPST catalyzes sulfate transfer from adenosine 3’-phosphate 5’-phosphosulfate (PAPS) to the hydroxyl group of specific tyrosyl residues (Stewart and Ronald, 2022). Both TPST and soluble sulfotransferase sequences contain a residue (Glu or His) whose side-chain provides the catalytic base, as well as PAPS-binding residues in two conserved motifs, the 5’ phospho-sulfate binding (5’-PSB) and the 3’ phosphate binding (3’-PB) elements (Stewart and Ronald, 2022). Although these conserved elements were not initially identified in the plant TPST sequence (Komori *et al*., 2009), recent analysis detected putative PAPS-binding motifs (Stewart and Ronald, 2022). AtTPST is a type I transmembrane protein, with a single membrane spanning segment and the carboxyl-terminus exposed to the cytosol. In contrast, animal TPST is a type II transmembrane protein, with the amino-terminus exposed to the cytosol. These different topologies and apparent absence of sequence similarity suggested that plant and animal TPSTs evolved independently via convergent evolution (Komori *et al*., 2009).

In Arabidopsis, TPST is expressed throughout the plant but most highly at the root apical meristem (Komori *et al*., 2009). An Arabidopsis *tpst* null mutant displays various developmental and growth phenotypes, including dwarfism and early senescence (Komori *et al*., 2009). TPST-mediated sulfation of tyrosine residues in PSYs, PSKs, RGFs, and CIFs is critical for high-affinity binding to their cognate receptors (Kaufmann and Sauter, 2019; Stewart and Ronald, 2022). The plant *TPST* gene is conserved from angiosperms to mosses (Zhao *et al*., 2019).

The spreading earth moss *Physcomitrium patens* is a tractable model for understanding early events in plant evolution (Rensing *et al*., 2020). Unlike angiosperms, whose genomes encode four classes of sTyr peptides (PSK, PSY, RGF, and CIF), which have multiple paralogs in each class, thus far, the *P. patens* genome has been found to only encode one copy of *PSY* and one copy of *PSK* (Furumizu *et al*., 2021). However, two immune-responsive PSY-like peptides in *P. patens* were recently identified (Lyapina *et al*., 2025). In addition to the ease of genome editing, *P. patens* is an advantageous model organism because it is predominantly haploid and undergoes expansion in two main ways. In juvenile stages, protonemal filaments expand via tip growth, and in adult tissues, gametophores form, in which the phyllids, or the leaf-like structure, expand using diffuse growth (Schumaker and Dietrich, 1998). By analyzing TPST function in *P. patens*, we can not only understand the role of sulfation in a system with fewer predicted sulfo-peptides, but also investigate both diffuse and tip growth. Thus, *P. patens* provides a platform from which to study plant TPST, such as identifying catalytic residues and defining substrate specificity determinants. Here we report that tyrosine-sulfated peptides are required for development, that the catalytic residue found in animal sulfotransferases is conserved in *P. patens*, and that PSY signaling-induced expansion is evolutionarily conserved.

## Results

### Sulfated peptides are required for development

To investigate the role of *TPST* (Pp3c16_24250) in cellular proliferation and development, we generated a null mutant using CRISPR/Cas9-mediated genome editing. *TPST* in *P. patens* has several predicted gene models, leading to two main predicted transcripts (Fig. S1A). The annotation in Phytozome suggests that both transcripts encode a single-pass type I transmembrane protein similar to Arabidopsis TPST (Komori *et al*., 2009; Goodstein *et al*., 2012; The UniProt Consortium, 2025). Additionally, both transcripts encode the same predicted signal peptide sequence. The only major difference between the two predicted proteins is the number of amino acids between the PAPs-binding catalytic region and the transmembrane domain (Fig. S1C). Both the short and long transcripts were detected using RT-PCR (Fig. S1B). Sequencing confirmed that the long transcript matches the predicted gene model V3.5, and the short transcript matches the predicted gene model V3.1. Therefore, we targeted a region of the locus corresponding to the amino-terminal coding region to create a *tpst* null mutant regardless of transcript length. Two independent *tpst* null mutant lines were recovered, each resulting from a frameshift near the protospacer cut site as a result of a deletion (Δ*tpst-6)* and an insertion (Δ*tpst-7)* (Fig. S1A). Both null mutants were smaller than wild-type plants (Fig. S1D and S1E). Δ*tpst-6* was selected for future study and is subsequently referred to as Δ*tpst*.

We found that Δ*tpst* plants had a dark center, appeared denser than wild type, and were very round (Fig. 1A). We measured total plant area and found that 14-day-old Δ*tpst* plants regenerated from protoplasts were 48% smaller than wild type (p < 0.0001) (Fig. 1B). To determine if the difference in plant area was a result of growth rate and or cell size, we measured chloronemal and caulonemal growth rates and cell size. While there was a trend that Δ*tpst* chloronemal and caulonemal apical cells grew slower, this was not statistically significant (Fig. S1F). We found that Δ*tpst* caulonemal subapical cells were 12% smaller than wild type (p = 0.0288), but there was no significant difference in chloronemal subapical cell length (Fig. S1G). Since the slightly shorter caulonemal cells do not account for the difference in total plant area, we quantified the number of caulonemal filaments emerging from the 14-day-old plants. We found that Δ*tpst* plants had 47% less caulonemal filaments compared to wild type (p < 0.001) (Fig. 1C), suggesting that the Δ*tpst* protonemal phenotype is a result of deficits of the transition from chloronema to caulonema. As an alternative method to measure the lack of caulonemal filaments, we quantified circularity, which is proportional to the ratio of the area to the square of the perimeter. The more caulonemata that extend from the plant, the lower the ratio. The difference in circularity closely reflects the reduction in the number of caulonemal filaments with wild-type plants exhibiting an average circularity of 0.023 and Δtpst 0.041 (Fig. 1D). As plants continued to mature, Δ*tpst* was unable to form expanded gametophores and instead appeared to abort growth once a bud was formed, despite forming the same amount of buds per area as wild type (Fig. S1H, Fig. 1E). Additionally, Δ*tpst* turned brown before wild type, an indication of early senescence (Fig. 1F). These data demonstrate that *TPST* function is critical for multiple aspects of *P. patens* development, including protonemal growth, gametophore expansion, and senescence.

**Figure 1.**
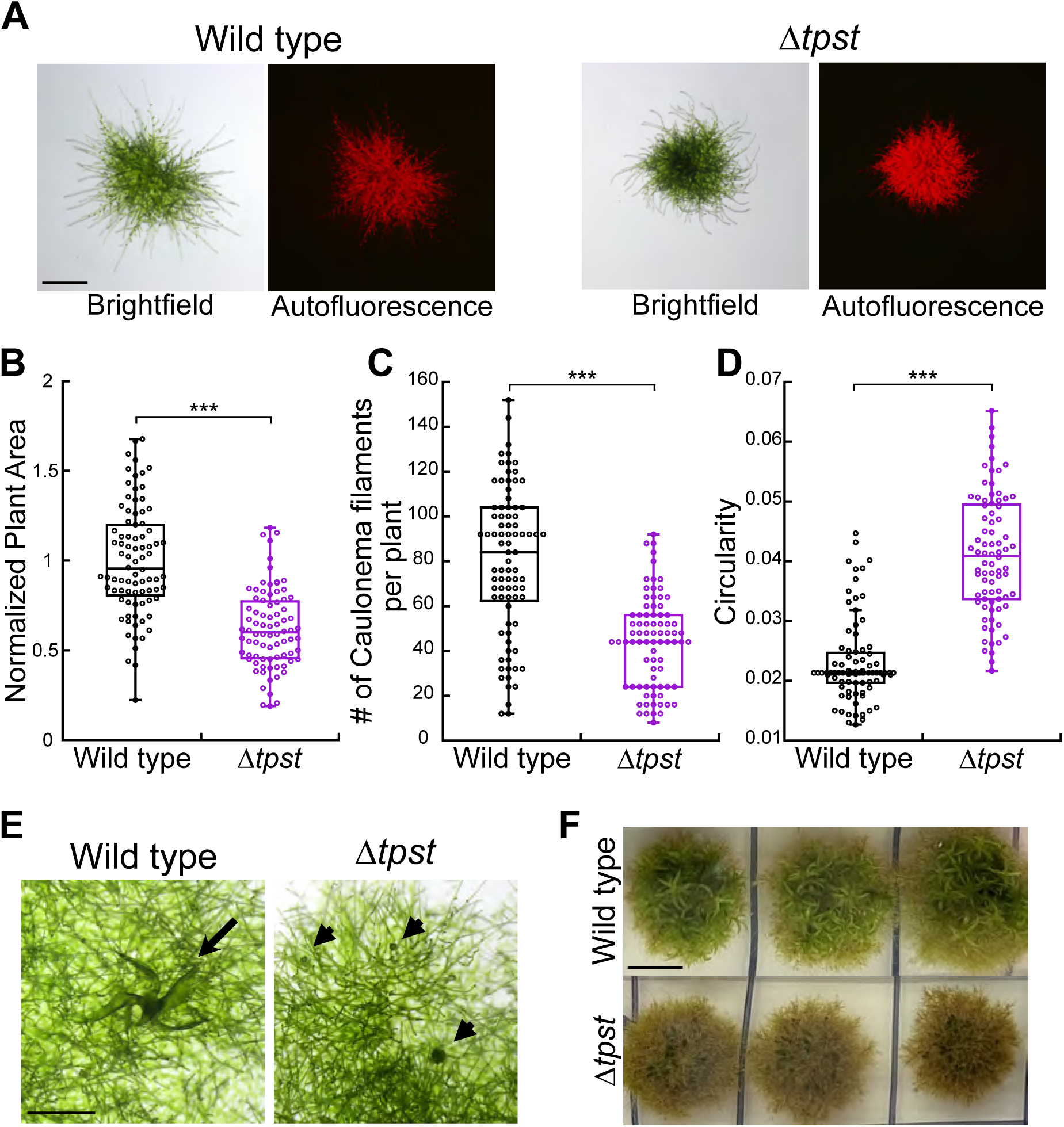
TPST is required for normal development. A) Representative brightfield and chlorophyll autofluorescence images of 2-week-old wild type and Δ*tpst* from plants that were regenerated from protoplasts on cellophane. Scale bar, 500 µm. B) Plant area was quantified and normalized to wild type. Experiments were repeated three independent times and the total number of plants quantified for each was: 84, WT; 80, Δ*tpst*. C) Number of caulonemal filaments protruding from the center per 2-week-old protoplast-regenerated plant. Filament number was calculated by counting the number per ¼ of plant and multiplying by 4. C) Quantification of circularity using the 2-week-old protoplast-regenerated plants measured in B. (B-D) Significance was calculated using a Kruskal-Wallis Test with a significance level of α=0.05. p<0.0001, for all comparisons. E) Representative brightfield images of 5-week-old tissue that had been regenerated from protoplasts. Arrow denotes a fully expanded gametophore. Arrowheads denote gametophore buds. Scale bar, 500 µm. F) 11-week-old whole plants that were regenerated from protoplasts. Scale bar, 0.5 cm.

### Sulfated peptides are required for normal cell division and expansion during gametophore development

To investigate why Δ*tpst* fails to form fully expanded gametophores, we used confocal microscopy to image bud development. We created a *tpst* null mutant in a moss line that stably expresses mEGFP-tubulin and has the plasma membrane-localized protein, TONNEAU1 (TON1), endogenously-tagged with 3XmRuby2 (Fig. S2A), which can be used to visualize the outline of cells during gametophore development (Wu *et al*., 2026). To follow bud development, we used live-cell imaging in microfluidic chambers (Bascom *et al*., 2016). The cell divisions underlying early bud formation in *P. patens* during gametophore development have been well-established (Harrison *et al*., 2009). The bud initial emerges similar to a protonemal side branch, but soon after emergence it appears swollen or bulbous in appearance (Harrison *et al*., 2009). It then undergoes an oblique cell division, followed by three subsequent asymmetric divisions, which establish a tetrahedral apical cell and mark the transition to three-dimensional growth (Harrison *et al*., 2009). Because of the marked appearance of a bud initial, the switch from filamentous to shoot fate is made prior to the first oblique division (Harrison *et al*., 2009). However, in Δ*tpst,* we noticed irregular bud morphology at the 4-cell stage. The bud initial appears bulbous, but the first division is not oblique, suggesting a delay in the decision between branch or bud fate (Fig. 2A). Time-lapse imaging beginning at the 6-cell stage showed that wild type undergoes a series of 12 cell divisions over the course of approximately 12 hours (Fig. 2B, Fig. S2B, Video 1). However, in the same time frame, Δ*tpst* underwent only 3 divisions (Fig. 2B, Fig. S2C, Video 2). As gametophore development progressed, Δ*tpst* continued to divide less than wild type and did not form a phyllid at the same time in development (Fig. 2C, Videos 3 and 4). Confocal imaging of early phyllid formation over the course of 23 hours showed that wild type performed both cellular expansion and division to form a phyllid (Fig. 2D, Video 5). However, a Δ*tpst* bud of approximately the same size (which is older than the wild-type bud because of fewer cell divisions) only expanded during the same 23-hour period (Fig. 2D, Video 6). Overall, defects in gametophore formation in Δ*tpst* were apparent from the onset of bud formation, characterized by deficits in cell division, including irregular first divisions and markedly slower rates of cell division.

**Figure 2.**
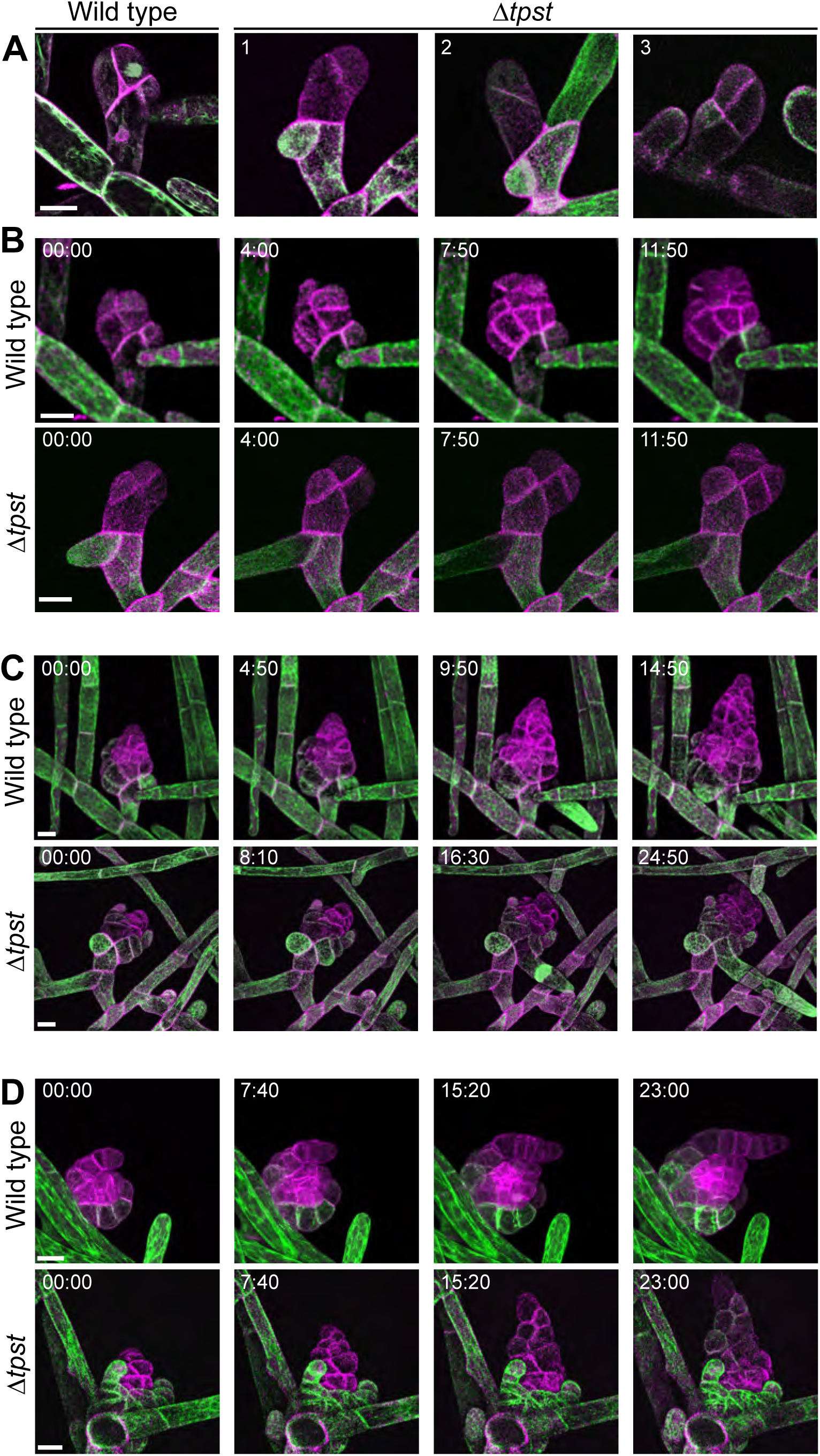
TPST is required for normal cell division during gametophore development. Plasma membranes are labeled with 3XmRuby2-TON1(magenta), and microtubules are labeled with mEGFP-tubulin (green). All images are maximum intensity projections of confocal Z-stacks. Scale bar for all images, 20 µm. A) Wild type and three example Δ*tpst* buds at the 4-cell stage. B) Time-lapse imaging starting at the 6-cell stage of the wild type bud from A and the Δ*tpst* bud labeled (1) from A. Also see Videos 1 and 2. C) Time-lapse imaging of the wild type and Δ*tpst* buds from B. Also see Videos 3 and 4. Time elapsed between acquisitions was approximately the same for both wild type and Δ*tpst.* D) Time-lapse imaging of phyllid expansion in a developing wild type and Δ*tpst* bud of approximately the same size. Also see Videos 5 and 6.

In addition to cell division, Δ*tpst* exhibited defects in cellular expansion during gametophore formation. During phyllid expansion, cells in a wild-type phyllid expanded as the leaf grew outwards, changing from a smaller, square shape to a rectangle, with an average of a 1.92-fold change in cell area over the course of 24 hours (Fig. 3A and B, Video 7). In contrast, cells in a Δ*tpst* phyllid retained an oblong morphology and did not significantly expand during a 24-hour imaging period, with an average increase in cellular area of only 1.08-fold (Fig. 3A and B, Video 8). Therefore, TPST function, and thus sulfated signaling peptides, are required for normal cellular expansion and division during gametophore development.

**Figure 3.**
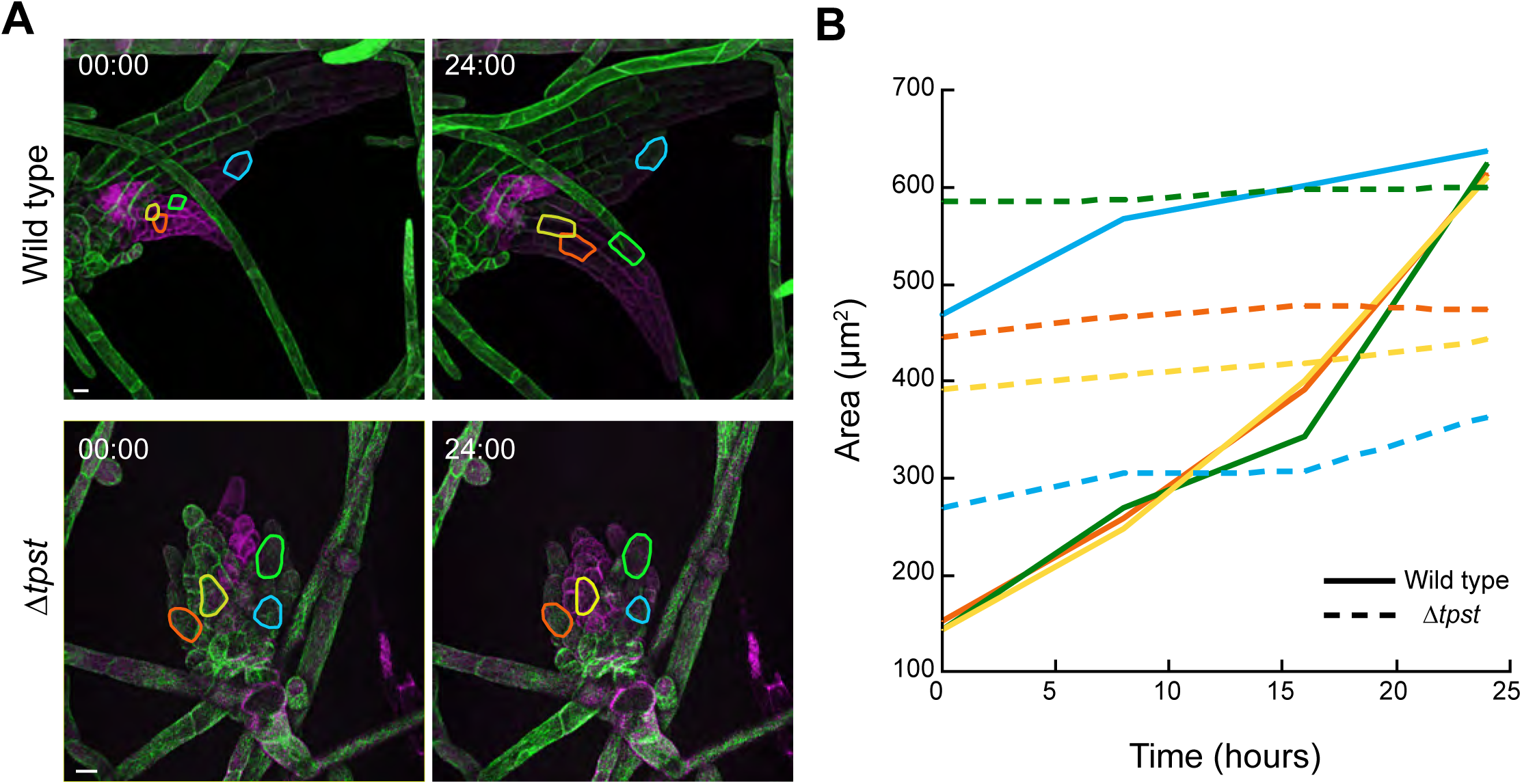
TPST is required for cellular expansion during gametophore development. (A) Time-lapse imaging of gametophore expansion in wild type and in Δ*tpst*. Plasma membranes are labeled with 3XmRuby2-TON1(magenta) and microtubules are labeled with mEGFP-tubulin (green). Orange, yellow, green, and blue lines indicate the outline of 4 cells across 24 hours. All images are maximum intensity projections of confocal Z-stacks. Scale bar for all images, 20 µm. B) Quantification of the change in area of the outlined cells in A across 24 hours. Colors on the graph correspond to the color outline in A. Solid lines indicate wild type. Dashed lines indicate Δ*tpst.* Also see Videos 7 and 8.

### Catalytic residues are conserved in PpTPST

Candidate 5’-PSB, 3’-PB, and catalytic base residues were identified in plant TPST sequences (Stewart and Ronald, 2022) (Fig. 4A). We therefore used CRISPR/Cas9 and homology-directed repair to generate Ala substitutions at three residues: Arg-81 in the putative 5’-PSB element, Arg-144 in the putative 3’-PB element, and His-124, the putative catalytic base (Fig. S3). Quantitative growth assays revealed that all three mutants (*tpst-R81A*, *tpst-H124A,* and *tpst-R144A*) were significantly smaller than the wild type (p < 0.001) but were not significantly different from the Δ*tpst* mutant or each other (Fig. 4C). This indicates that all three mutants had compromised TPST activity. However, there were differences in protonemal morphology amongst the mutants. The *tpst-H124A* mutant was dense, round, and had fewer caulonemal filaments, similar to the Δ*tpst* mutant (Fig. 4B). In contrast, the *tpst-R81A* and tpst-*R144A* mutants were less dense and exhibited more caulonemal filaments.

**Figure 4.**
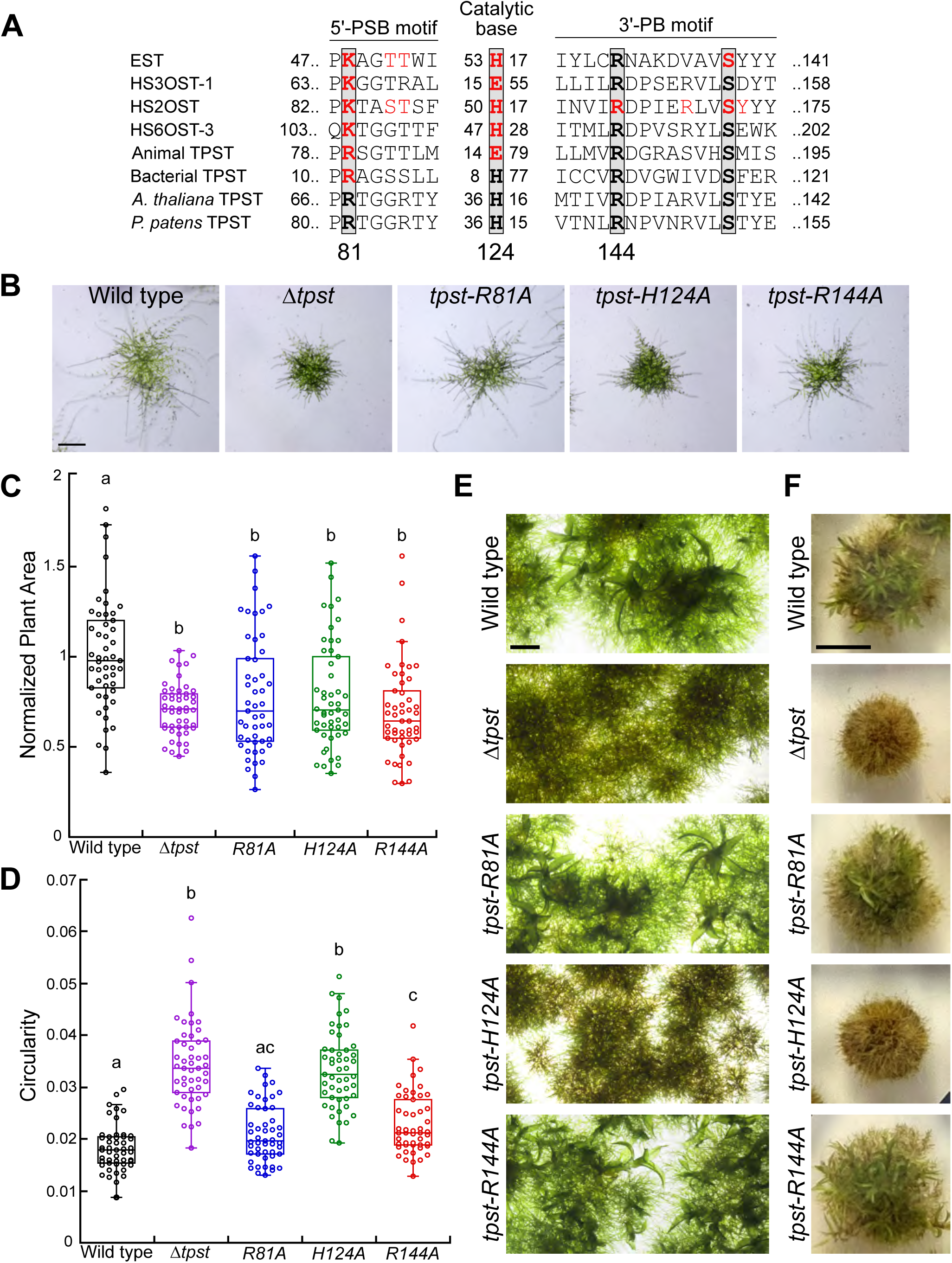
Histidine 124 is catalytically important for TPST function. A) Sequence motifs of different sulfotransferases across organisms showing the 5’-PSB, candidate catalytic base, and 3’-PB. Adapted from Stewart and Ronald, 2022 to include *P. patens* TPST. Numbers on the left and right of the motifs indicate the starting and ending position. The numbers between motifs indicate the number of residues in between the selected sequences. Substitutions of residues in red decrease enzymatic activity. Numbers underneath the black boxes indicate the position in *P. patens* TPST. Enzymes are EST, estrogen 17-β sulfotransferase (*Mus musculus*; Protein Data Bank [PDB] code: 1AQU); HS3OST-1, heparan sulfate (3-O) sulfotransferase (*M. musculus*; PDB code: 1S6T); HS2OST, heparan sulfate (2-O) sulfotransferase (*Homo sapiens*; PDB code: 3F5F); HS6OST-3, heparan sulfate (6-O) sulfotransferase (*Danio rerio*; PDB code: 5T03); animal TPST (*H. sapiens*; PDB code: 5WRI); bacterial TPST (*Xanthomonas spp*.; GenBank accession number: WP_027703307); *A. thaliana* TPST (*Arabidopsis thaliana*; BAI22702); *P. patens* TPST (NCBI Reference Sequence: XM_024542583.2). B) Representative brightfield images of 2-week-old plants regenerated from protoplasts on cellophane. Scale bar, 500 µm. C and D) Quantification of plant area and circularity using images of 2-week-old regenerated protoplasts. Plant area is quantified and normalized to wild type. Significance was calculated using a Kruskal-Wallis Test with a significance level of α=0.05. Experiments were repeated two independent times, and a total number of 49 plants was quantified for each line. E) Representative brightfield images of 3-week-old tissue on cellophane. Scale bar, 500 µm. F) 8-week-old whole plants from protoplast-regenerated tissue. Scale bar, 0.5 cm.

To quantify these differences in plant morphology, we measured the circularity of each plant as a readout for the amount of caulonemal filaments. Wild-type plants had an average circularity of 0.0182. Δ*tpst* and *tpst-H124A* plants had significantly higher circularity values of 0.0344 and 0.0333, respectively, whereas *tpst-R81A* (0.0210) and *tpst-R144A* (0.229) displayed intermediate values (Fig. 4D). These results demonstrated that the *tpst-H124A* alteration phenocopied the defect in transition to caulonemal filaments associated with the Δ*tpst* null allele, whereas the *tpst-R81A* and *tpst-R144A* alterations conferred an intermediate phenotype.

To assess whether these alterations affect gametophore development or senescence, we continued to monitor the growth and development of the mutants. After 8 weeks, the Δ*tpst* and *tpst-H124A* mutants exhibited early senescence in comparison to the wild type, the *tpst-R81A* and *tpst-R144A* mutants (Fig. 4F). Additionally, the *tpst-H124A* mutant, like the Δ*tpst* mutant, did not form expanded gametophores, whereas *tpst-R81A* and *tpst-R144A* plants formed fully expanded gametophores (Fig. 4E). Together, the decreased average plant area combined with intermediate circularity values despite forming fully expanded gametophores suggest that the R81A and R144A alterations have lost some TPST function. In contrast, the H124A and Δ*tpst* alterations are indistinguishable, indicating that residue His-124 is essential for TPST function.

### PSY is evolutionarily conserved

Because Δ*tpst* plants presumably cannot produce tyrosine sulfated peptides (Komori *et al*., 2009; Matsuzaki *et al*., 2010; Kaufmann, Stührwohldt and Sauter, 2021), we investigated whether exogenous application of *P. patens* PSY1 (PpPSY1), *A. thaliana* PSY1 (AtPSY1), and *A. thaliana* PSK1(AtPSK1) synthetic peptides could alter the Δ*tpst* phenotype. Initially, we exposed 8-day-old tissue regenerated from protoplasts to peptides for 12 days and screened for gametophore formation. The Δ*tpst* mutant formed expanded gametophores after exposure to 200 nM of AtPSY1 or 1 µM of PpPSY1, but not to 200 nM of AtPSK1 (Fig. 5A). We repeated this experiment to test several doses of peptide to determine the optimal concentration (Fig. 5B). To observe expanded gametophores in Δ*tpst*, we used a higher concentration of PpPSY1 compared to AtPSY1.

**Figure 5.**
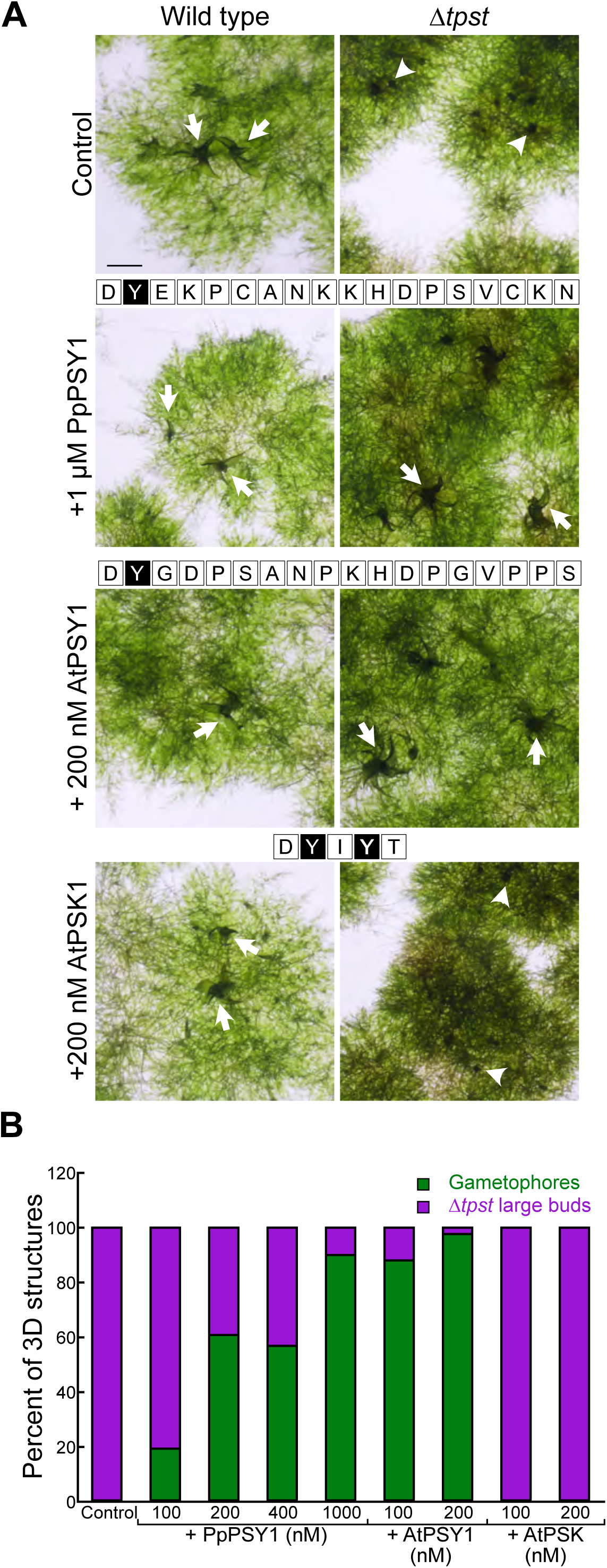
Addition of exogenous PSY1 induces gametophore expansion in Δ*tpst*. A) 20-day-old protoplast-regenerated plants on cellophane after 12 days on PpNH4 media supplemented with either 1 µm PpPSY1, 200 nM AtPSY1, or 200 nM AtPSK1. Arrows denote an example of an expanded gametophore. Arrowheads denote an example of a gametophore bud. The 18 amino acid sequence of PSY1 in *P. patens* (moss) and *A. thaliana,* and the 5 amino acid sequence of PSK1 in *A. thaliana* are shown above the corresponding image. Black boxes represent the sulfated tyrosine residues. Scale bar, 0.1 cm. B) Quantification of gametophore formation compared to Δ*tpst* bud formation on approximately 3-week-old protoplast-regenerated Δ*tpst* plants grown on media supplemented with varying doses of peptide.

To quantify the peptide-mediated rescue of protonemal growth in Δ*tpst*, we moved 7-day-old plants regenerated from protoplasts to media supplemented with AtPSY1 or PpPSY1. After 5 days of peptide treatment, Δ*tpst* plants closely resembled wild type (Fig. 6A). Interestingly, wild-type plants grown on either peptide were significantly larger than wild-type plants in the absence of peptide (Fig. 6B). There was no significant difference between the Δ*tpst* and wild-type plants grown on AtPSY1 (p > 0.05). When grown on PpPSY1, Δ*tpst* plants were the same size as the wild-type controls. However, in comparison to all other peptide conditions, Δ*tpst* grown on PpPSY1 were the smallest. These observations support that the PpPSY1 peptide is less effective at rescuing Δ*tpst* than AtPSY1. However, both PpPSY1 and AtPSY1 in the media were able to rescue the circularity phenotype of Δ*tpst*, indicating that the addition of sulfated peptide rescues the developmental defect in the transition from chloronema to caulonema (Fig. 6C).

**Figure 6.**
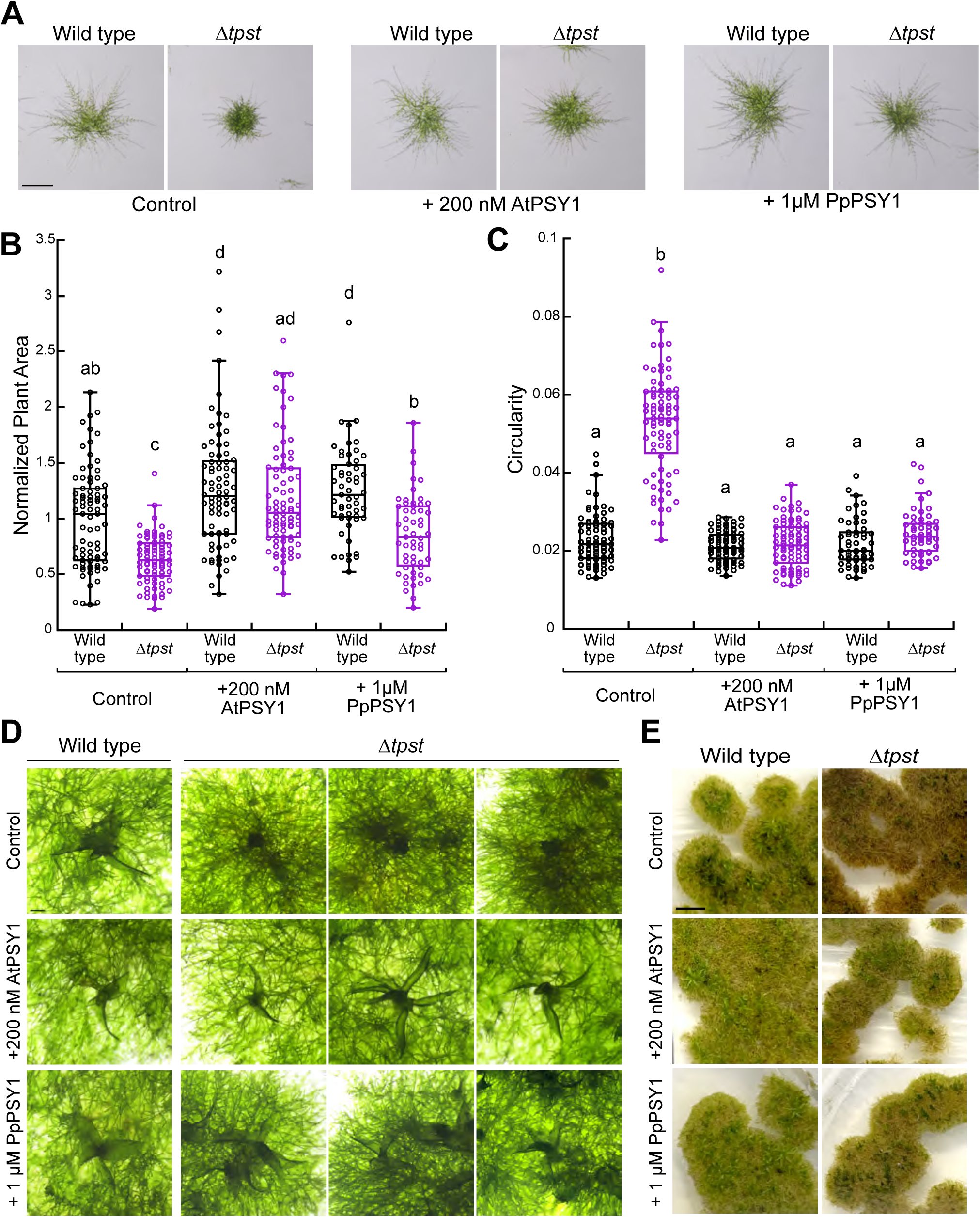
Exogenous PSY1 enhances growth. A) Representative brightfield images of wild type and Δ*tpst* from 12-day-old regenerated protoplasts on cellophane. Scale bar, 500 µm. B and C) Quantification of plant area (B) and circularity (C) using images of 12-day-old regenerated protoplasts on cellophane. All values are normalized to the wild-type control. Letters indicate groups that are significantly different as determined by a Kruskal-Wallis test (α = 0.05) with a Dunn’s post hoc test and Bonferroni correction. Experiments repeated three independent times and the total number of plants quantified for each was: 80, WT and Δ*tpst* control; 80, WT and Δ*tpst* + 200 µM AtPSY1; 55, WT and Δ*tpst* + 1µM PpPSY1. D) Representative brightfield images of expanded gametophores in 20-day-old protoplast-regenerated wild type and Δ*tpst* plants on cellophane after 15 days on peptide media. Scale bar, 200 µm. E) 7-week-old whole plants from protoplast-regenerated tissue on cellophane. Scale bar, 0.5 cm.

We continued to transfer the protoplast-regenerated plants on cellophane to fresh peptide media every week and confirmed our previous results that in the presence of either peptide, Δ*tpst* exhibited expanded gametophores (Fig. 6D). There was no noticeable difference in wild-type gametophore expansion in the presence of AtPSY1 or PpPSY1. As the protoplast-regenerated plants continued to age, we found that Δ*tpst* turned brown around 7 weeks after protoplasting (tissue that is grown on cellophane behaved differently than tissue that was moved directly onto media). However, Δ*tpst* grown on peptide media remained green, similar to wild type, suggesting that in the presence of the peptide, Δ*tpst* did not age as quickly as Δ*tpst* under control conditions (Fig. 6E). Δ*tpst* grown on either PpPSY1 or AtPSY1 did not senesce at the same time as Δ*tpst* under control conditions. Therefore, supplementing media with exogenous PSY1 rescued all developmental aspects of the Δ*tpst* phenotype, restoring normal protonemata, gametophore expansion, and timing of senescence. Notably, the ability of AtPSY1 to rescue Δ*tpst* in *P. patens* suggests that the PSY signaling pathway is evolutionarily conserved.

Given that Arabidopsis PSY1 peptide influences *P. patens* development, we tested whether PSY from *P. patens* affects other plant species. Previous work demonstrated that exogenous application of AtPSY1 and OsPSY1 peptides promote root growth in Arabidopsis and rice (Pruitt *et al*., 2017; Ercoli *et al*., 2024; Shi *et al*., 2024), so we conducted root assays in these species using the synthetic PpPSY1 peptide. In Arabidopsis, we were able to demonstrate that both wild-type (Col-0) and *tpst-1* plants showed an increase in root elongation in response to synthetic AtPSY1 and PpPSY1 treatment, when compared to mock-treated plants (Fig. 7 A,B). However, the impact of PpPSY1 on wild-type Arabidopsis roots is stronger at 250nM (p = 0.0034) compared to 100nM (Fig. 7B, left panel). Additionally, exogenous application of PpPSY1 peptide promoted root elongation in wild-type Kitaake seedlings (Fig. 7C, D). We repeated this experiment with the addition of the synthetic AtPSY1 and OsPSY1 peptides (Fig. S4); results show that PSY peptides from moss, Arabidopsis, and rice promote root elongation of Kitaake seedlings (p < 0.0005) (Fig. S4). These results further support that PSY signaling is evolutionarily conserved.

**Figure 7.**
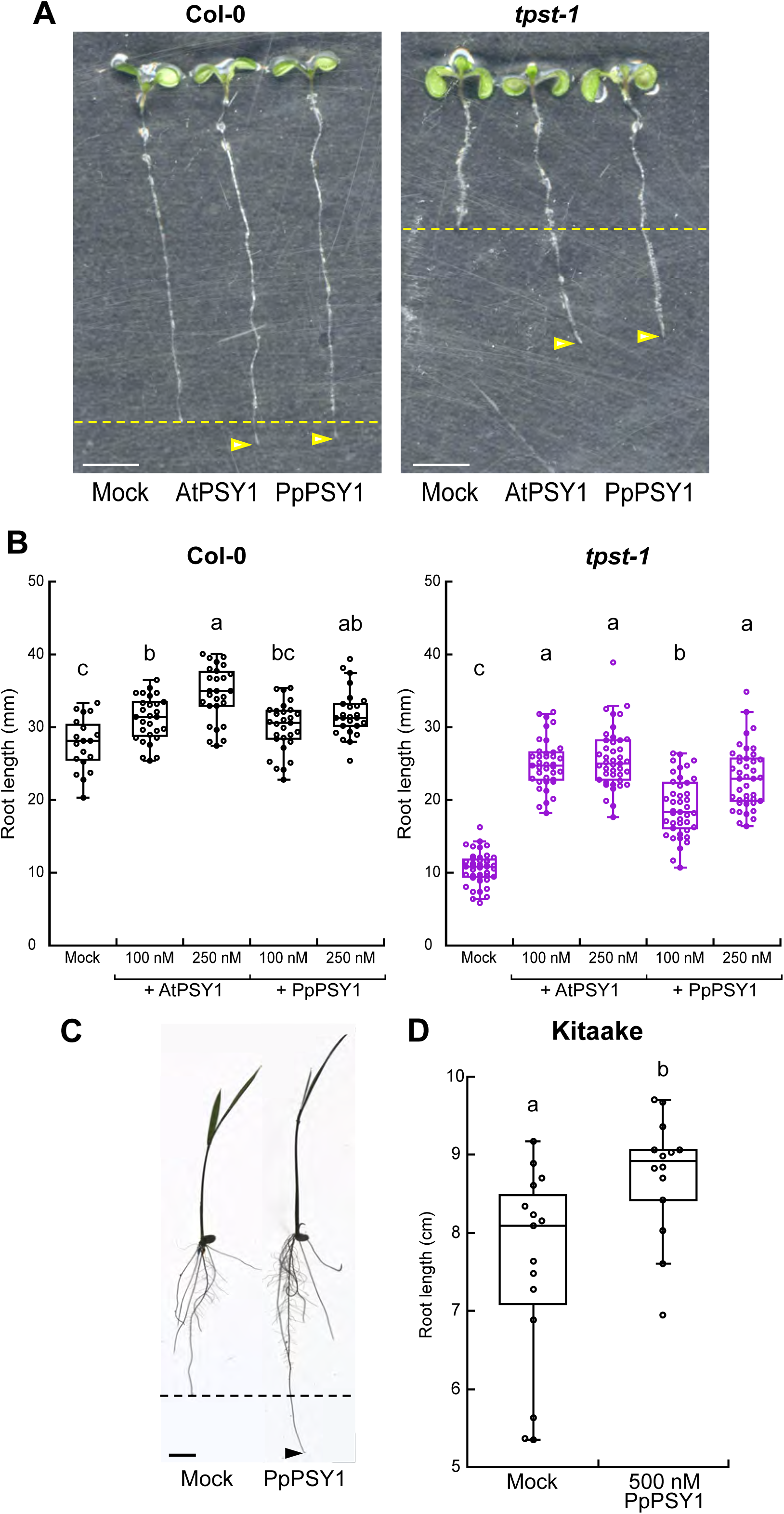
Synthetic PpPSY1 peptide promotes root elongation in Arabidopsis & Rice seedlings. (A) Representative images of primary root growth of Col-0 (wild type, left) or *tpst-1* (right) seedlings 6-days-post germination (dpg) grown on 1X MS vertical plates with or without synthetic peptide treatment (250nM AtPSY1 or PpPSY1). Enhanced root elongation of peptide treated samples indicated by the yellow serrated line, with the arrows indicating the primary root tip. Scale represents 5mm. (B) Quantification of primary root elongation for Col-0 (left, n=19-29) or *tpst-1* (right, n=42-44) seedlings grown on 1X MS vertical plates with or without synthetic peptide PpPSY1 or AtPSY1 treatment (100nM or 250nM). 6-dpg for Col-0 and 8-dpg for *tpst-1*. Different letters indicate significant differences, as determined by a Kruskal-Wallis test (α = 0.05) with a Dunn’s post hoc test and Bonferroni correction. Experiments repeated three independent times. (C) Representative image of primary root growth of Kitaake (wild type) seedlings grown hydroponically in 1X MS with or without synthetic PpPSY1 treatment (500nM), 6-days-post treatment. Enhanced root elongation of peptide treated samples indicated by the black serrated line, with the arrows indicating the primary root tip. Scale bar represents 1cm. (D) Quantification of root elongation of Kitaake seedlings (n= 14-15), 6-dpg, grown on 1X MS plates with or without synthetic 500nM PpPSY1 treatment. Different letters indicate significant differences, as determined by a Kruskal-Wallis test (α = 0.05). Experiments repeated three independent times.

## Discussion

Here, we report that sulfotyrosyl peptides are critical for cellular expansion and division, as Pp*tpst* null mutants (Δ*tpst*), which are deficient in producing sulfated peptides, have defects in cell expansion in the juvenile (protonemata) and adult (gametophores) phases of moss growth. During gametophore development, Δ*tpst* exhibited an abnormal first division, and decreased rates of cell division and cellular expansion. Additionally, Δ*tpst* plants age earlier than wild-type plants.

The Δ*tpst* developmental phenotypes in *P. patens* were largely rescued by exogenous application of AtPSY1. Reciprocally, exogenous addition of PpPSY1 to both Arabidopsis and rice elicited root elongation. These results indicate that PSY peptides act as functional orthologs in plant development and further suggest that peptide recognition may function through an evolutionarily conserved signaling pathway. While PSY receptors have been identified and characterized in Arabidopsis (Ogawa-Ohnishi *et al*., 2022), future work will aim to elucidate the receptors in *P. patens* and their downstream signaling modules. The comparison of peptide function in different plant species and the binding of orthologous receptors will shed light on the conserved biological functions mediated by PSY-signaling.

Overall, our findings indicate that PSY signaling is conserved in *P. patens* and is critical for development. Null mutants of *TPST* in both Arabidopsis (*tpst-1*) and *P. patens* (Δ*tpst)* exhibit dwarfism and decreased plant area, respectively, and display early senescence. These phenotypic similarities suggest possible functional conservation of TPST. The lower efficacy of PpPSY1 in rescuing Δ*tpst and promoting root elongation in Arabidopsis and rice* might be attributed to the PpPSY1 peptide sequence containing two cysteine residues, which are not a conserved feature of PSY peptides (Amano et al., 2007; Kaufmann and Sauter 2019), and give the synthesized peptide the potential to be easily oxidized over time. The inability of AtPSK1 to rescue Δ*tpst* might be explained by the possibility that PSK1 may not be sulfated in *P. patens*, and thus not dependent on TPST function. The putative PSK1 sequence from *P. patens* (Furumizu *et al*., 2021) lacks the highly conserved aspartic acid residue that precedes the PSK1 peptide sequence. All known sulfated peptides, with the exception of GLV9, contain a DY motif (Kaufmann and Sauter, 2019), which is absent from the PSK1 sequence in *P. patens*. The aspartic acid residue that precedes the sulfated tyrosine residue is one of the most important factors for directing tyrosine sulfation in PSK1 (Matsubayashi and Sakagami, 1996). Given the absence of this motif from PpPSK1, which instead has an FY motif, it is possible that the mature peptide lacks a sulfated tyrosine, suggesting that PSK signaling in *P. patens* may not require tyrosine sulfation.

Recently, two immune-responsive small secreted peptides were identified in *P. patens* as sharing similarities with the PSY peptide motif, creating a new category of sulfated peptides, PSY-like (PSYL) (Lyapina *et al*., 2025). Quantification of protonemal cell elongation of wild-type plants on media containing varying concentrations of both the sulfated and non-sulfated versions of PSY, PSYL1, and PSYL2 showed that the sulfation of the tyrosine residue is necessary for activity of the PSY and PSYL peptides in *P. patens.* This result is consistent with our finding that TPST function and, therefore, its post-translational modification of sTyr peptides, were critical for proper development. The addition of sulfated PSY, PSYL1, or PSYL2 to the media increases cell length, which is in line with our findings that the addition of exogenous PSY increased plant area. Both PSYL1 and PSYL2 peptide sequences contain a motif that resembles the PSY peptide motif (DY*(D)(P)**N**H*P), but the conserved His and Pro residues are in different positions than in PSY (Lyapina *et al*., 2025). Given the resemblance to PSY, it is possible that the addition of exogenous PSYL1 and PSYL2 could rescue some of the developmental deficits of Δ*tpst*, including protonemal plant size, gametophore expansion, and senescence.

Moss mutants carrying an alanine substitution of the candidate catalytic base, His-124 in PpTPST, resembled the moss Δ*tpst* mutant phenotype. This indicates that His-124 is essential for protein function and thus necessary for proper plant development. These observations agree with prior findings that the histidine in the active site of *M. musculus* EST and *F. chlorifoli*a F3ST is required for enzymatic activity (Marsolais and Varin, 1997; Kakuta *et al*., 1998). Thus, the conserved histidine found in the active site of other studied sulfotransferases, which is required for catalytic activity, is conserved in plant TPSTs. Based on this finding and the identification of putative PAPs-binding motifs in plant TPSTs (Stewart and Ronald, 2022), it is possible that plant and animal TPSTs did not evolve via convergent evolution as first suggested (Komori *et al*., 2009).

Although *tpst-H124A* most closely resembled Δ*tpst*, the other point mutants, *tpst-R81A,* and *tpst-R144A*, are the same size as Δ*tpst*, but with a lower circularity, which could potentially be attributed to the point mutants retaining some weak PAPS binding. Prior studies have found that some missense substitutions in the 5’-PSB and 3’-PSB motifs of other sulfotransferases retain some wild-type enzymatic activity (Bick *et al*., 2010; Liu and Pedersen, 2022). Therefore, this residual activity could help explain the small protonemal phenotype of *tpst-R81A* and *tpst-R144A*. This finding suggests that plant TPSTs use the same catalytic mechanism as animal sulfotransferases.

## Materials and Methods

### Tissue propagation

*P. patens* lines were propagated weekly by moderate homogenization in water and then plated onto PpNH4 (1.03 mM MgSO4, 1.86 mM KH_2_PO_4_, 3.3 mM Ca(NO_3_)_2_, 2.7 mM (NH_4_)_2_-tartrate, 45 μM FeSO_4_, 9.93 μM H_3_BO_3_, 220 nM CuSO_4_, 1.966 μM MnCl_2_, 231 nM CoCl_2_, 191 nM ZnSO_4_, 169 nM KI, and 103 nM Na_2_MoO_4_, containing 0.7% agar) plates covered with cellophane (Wu and Bezanilla, 2014). Tissue was grown in Percival growth chambers at room temperature under 85 µmol_photons_/m^2^s light with long-day (16/8) conditions. Peptide media was created by pouring 25 mL PpNH4 plates supplemented with either 200 nm AtPSY1, 200nM AtPSK1, or 1 µM PpPSY1.

### RT-PCR

We isolated total RNA from 8-day old wild-type plants regenerated from protoplasts using an RNeasy Plant mini prep kit (Qiagen). cDNA was prepared from 1µg of total RNA using a SuperScript III reverse transcriptase kit, following the manufacturer’s recommendation (Invitrogen). We amplified the TPST transcripts using the primers listed in Supplementary Table 2 using 1 µL of cDNA.

### Generation of knockout and point mutation lines

Approximately one-week-old tissue was used for transformations. Transformations into *P. patens* protoplasts were done using the PEG-mediated transformation protocol described in (Wu, Ryken and Bezanilla, 2023). To generate knockouts (Δ*tpst-6)*, 15 µg of the CRISPR plasmid (pMH-Cas9-sgRNA plasmid backbone) was used. For HDR transformations using annealed oligonucleotides (Δ*tpst-7, tpst-R81A, tpst-H124A, tpst-R144A*), 15 µg of the CRISPR plasmid and 10 µL of annealed oligonucleotides were used. The sequences of each oligonucleotide and protospacer used are located in Supplementary Table 2.

After transformation, protoplasts were plated onto PRMB media covered in cellophane and left to recover for 4 days. The cellophane from the PRMB media (PpNH4 medium supplemented with 8.5% mannitol and 10 mM CaCl_2_), which contains the protoplasts, was moved to selection (PpNH4 media containing 15 µg ml^-1^ hygromycin) for 1 week. Surviving protoplasts have taken up the CRISPR plasmid. After 1 week of selection, the cellophane was moved to PpNH4 media. After 1-2 weeks, protoplast-regenerated plants were individually picked to fresh PpNH4 plates without cellophane, before being genotyped around 2 weeks later.

### Genotyping

Genomic DNA was extracted from *P. patens* tissue (Wu, Ryken and Bezanilla, 2023). Competition PCR was used to genotype plants. All genotyping primers are listed in Supplementary Table 2. Competition PCR uses two primers, one forward and one reverse, that are around 500 base pairs upstream and downstream of the protospacer region. In addition, a third internal primer was used, which was designed to anneal to the CRISPR cut site. Therefore, in edited plants, the internal primer does not bind, and only the larger PCR product is obtained. If the plant is not edited, the shorter PCR product, using the internal primer, is preferentially produced. Any plants that had a larger PCR product were used to PCR amplify the targeted region a second time, using only the external primers. The PCR product was sequenced using Sanger sequencing.

To identify point mutations, a second competition PCR was conducted using the same external primers and one primer that annealed to the desired point mutation and silent mutations. Therefore, in plants with the correct point mutation, the internal primer binds, forming the smaller PCR product. Any plants without the correct point mutation had a larger PCR product, as the internal primer could not bind. DNA from plants producing the smaller product was then amplified again with the only external primers, and those products were sequenced using Sanger sequencing.

### Imaging and morphometric analysis of growth assays

Protoplast preparation for growth assays was performed using a standard protocol (Wu, Ryken and Bezanilla, 2023). Approximately one week old ground tissue was protoplasted and plated onto PRMB media for 4 days. Afterwards, the protoplasts on cellophane were moved to PpNH4 media. All growth assays were performed using 2-week-old plants except for the data presented in Fig. 6.

Brightfield and chlorophyll autofluorescence images were collected using a stereomicroscope (Nikon SMZ25) equipped with a CCD color camera (Nikon digital DS-Fi2) with a 1X objective. Acquisition settings were held at the same settings for each image. Chlorophyll autofluorescence was captured using a 480/40 excitation filter with a 510 long pass emission filter. The red channel of the fluorescence image corresponds to chloroplast autofluorescence.

To ensure image acquisition occurred with the sample identity blinded to the observer, the genotype of each line was hidden at the time of imaging. Each line was imaged on the same day, and 30 images of individual plants were taken from each line. Both brightfield and fluorescence images were collected. To quantify plant area, the red channel was separated from each RGB fluorescence image. Images from the red channel were then analyzed using a macro in ImageJ (Galotto, Bibeau and Vidali, 2019). The threshold was set at max entropy and was manually set for each plant. The total area of each plant was calculated using the largest thresholded object in a cropped window.

For growth assays pertaining to plants grown on peptide, 1-week-old protoplasts on cellophane were transferred to peptide media or a fresh plate of PpNH4 as the control. After 5 days on peptide, 12-day-old protoplast-regenerated plants were imaged for growth assays. Growth assays were analyzed as described above. The cellophanes containing the protoplast-regenerated plants were moved to freshly poured peptide media or PpNH4 for the control on a weekly basis. Plants were moved to fresh media each week until senescence occurred.

### Senescence

To check whether mutant plants have a senescence phenotype, individual plants from protoplast-regenerated tissue were placed on the same plate of PpNH4 and left in the growth chamber for several weeks until at least one line turned brown (senescence). Senescence images were taken using a cellphone.

### Brightfield microscopy

For protonemal growth rate and subapical cell length, 2-week-old protoplast-regenerated plants were mounted on a square agar pad on a glass slide and submerged in Hoagland’s medium (4 mM KNO_3_, 2 mM KH_2_PO_4_, 1 mM Ca(NO_3_)_2_, 89 µM Fe citrate, 300 µM MgSO_4_, 9.93 µM H_3_BO_3_, 220 nM CuSO_4_, 1.966 µM MnCl_2_, 231 nM CoCl_2_, 191 nM ZnSO_4_, 169 nM KI, 103 nM Na_2_MoO_4_) (Wu and Bezanilla, 2014). The coverslip was sealed with VALAP (1:1:1 parts of Vaseline, lanoline, and paraffin). Images were acquired using a Nikon Ti microscope equipped with a 0.8 NA 20X objective. Protonemal growth rate was calculated by measuring the change in length of a growing tip cell during a 2-hour time-lapse using ImageJ. Multiple XY positions were acquired at a single focal plane. White light was continuously provided throughout the time-lapse to support plant growth. Subapical cell length was determined by measuring the distance between subapical cell plates in ImageJ.

### Laser scanning confocal microscopy

Homogenized plant tissue was pipetted into a PDMS microfluidic chamber (Bascom *et al*., 2016), followed by an infusion of Hoagland’s medium. The chamber was then submerged in Hoagland’s medium under 85 µmol_photons_/m^2^s light with long-day (16/8) conditions until plants were ready to image (Wu *et al*., 2023). Confocal images were captured using a Nikon A1R laser scanning confocal with a 0.75 NA 20X objective (Nikon) at room temperature. Acquisition was controlled by NIS-Elements. Laser illumination was set to nm for 3XmRuby (laser power, 1%; Gain, 35-50; PMT offset 20-28). Emission filter was 595/50 nm and collected with a GaAsp PMT. Images were deconvolved using 20 iterations of Richardson-Lucy on NIS-Elements software. For time-lapses, multiple XY positions were acquired at several focal planes. Images were acquired every 10 minutes. White light was provided between acquisition times to support plant growth.

### Bud and gametophore quantification

Images were collected using a stereomicroscope (Nikon SMZ25) equipped with a CCD color camera (Nikon digital DS-Fi2) with a 1X objective. For quantification of bud and gametophore formation of wild type versus Δ*tpst* (Fig. S1H), images of 14-day-old ground tissue were collected. The number of buds and gametophores per field of view (32.0972763 µm^2^) was counted and averaged. For quantification of gametophore formation under various peptide conditions, images were acquired of approximately 3-week-old protoplast-regenerated Δ*tpst* plants. The number of gametophores or Δ*tpst* buds was counted for each image. The proportion of gametophores or Δ*tpst* buds per condition was calculated.

### Cellular expansion quantification

24 hour time-lapses were acquired using maximum intensity projections of confocal images. 4 cells in wild type and Δ*tpst* were identified and outlined using the polygon tool in ImageJ. The area of each polygon was measured. The same cell was identified at the 8-hour, 16-hour, and 24-hour mark. The area was measured for each cell at each time point.

### Peptide Synthesis

The peptides used in this study include PpPSY1 (DY^S^EKPCANKKHDPSVCKNG), AtPSY1 (DY^S^GDPSANPKHDPGVPPSA), OsPSY1 (DY^S^PAPGANPRHNPKRPPG), and AtPSK1 (Y^S^IY^S^TQ). All peptides are tyrosine-sulfated as indicated (Y^S^). The synthetic AtPSY1 peptide used in these experiments lacks the hydroxy- and L-Ara3-modifications at the C-terminus. All peptides were obtained from Pacific Immunology (Ramona, CA, USA) and resuspended in double distilled water.

### Root Growth Assays

*Arabidopsis thaliana* ecotype Columbia (Col-0, wild type) and *tpst-1* (SALK_009847) seeds were surface sterilized with 70% ethanol for 10 minutes, rinsed three times with 100% ethanol, and then stratified in 0.1% agarose for 2-4 days at 4°C before germination. Seedlings were sown on standard 1X MS media plates (Murashige Skoog salt mixture with vitamins, MSP09-Caisson Laboratories), 1% sucrose, and 0.3% gellan gum (G024-Caisson-Gelzan), adjusted to pH 5.8 with KOH. MS plates were supplemented with various concentrations of synthetic PpPSY1, AtPSY1, or water (mock control). For each experiment, MS media was freshly prepared and cooled in a 55-60°C water bath after autoclaving before adding chemicals. Seeds were placed on the plates (15-20 seeds per plate) and the lids were secured with Micropore surgical tape (1530-0). For root assays, plants were grown in vertically positioned plates in a chamber under long day conditions (16 hours light:8 hours dark) at 21°C. Germination for each seedling was marked, and primary root length was marked every day until the end of the experiment (6-or 8-days post germination, dpg). Arabidopsis root plates were scanned and primary root growth was measured using Fiji Is Just ImageJ (Version 2.16) (Schindelin *et al*., 2012).

*Oryza sativa* ssp. *Japonica* cultivar Kitaake (wild type) seeds were dehusked, surface-sterilized in 20% bleach for 30 minutes, and thoroughly washed with autoclaved water. The seeds were grown on 1X MS media (MSP09-Caisson Laboratories) with 1% sucrose (pH 5.8), either as solid media plates containing 0.3% gellan gum or hydroponically in flasks with 100 mL of 1X MS without gellan gum. For solid medium conditions, seeds were sown on plates (15 per plate), sealed with Micropore tape, and placed vertically for 7 days. For hydroponic conditions, flasks containing 1X MS were cooled before adding peptides and sterilized seeds (15 per flask), and then placed in the rice incubator for 6 days. The incubators used for rice germination and growth were set with a 14-h-light/10-h-dark photoperiod at 28°C/24°C. MS plates or flasks were supplemented with synthetic PpPSY1, AtPSY1, OsPSY1, or water (mock control); the concentration of the peptide treatment is specified in the figure legends. For rice seedlings grown on plates, the germination was marked, and the root length was marked every day until the end of the experiment (6 dpg). Rice seedlings grown hydroponically or on solid media were imaged, and root length measurements were done in Fiji Is Just ImageJ (Version 2.16) (Schindelin *et al*., 2012).

### Statistical Analyses

A non-parametric approach was taken to analyze differences between different experimental groups. Statistical significance was determined using a Kruskal-Wallis test followed by a post-hoc Dunn’s test when a significant difference was found. A Bonferroni correction was applied to the p-values resulting from Dunn’s test to control for multiple comparisons. Pairwise adjusted p-values at a significance level of α = 0.05 were used to identify groups that were statistically different from each other. The compact letter display (CLD) approach was used to categorize each group into statistically distinguishable subsets. Letters were assigned to indicate significance between groups. Groups with the same letter were not statistically different. Groups with non-overlapping letters are not statistically different. Analyses were performed using R version 4.4.1 (Team, R. C., 2024).

## Acknowledgements

We thank Maria Florencia Ercoli for the Arabidopsis *tpst-1* seeds and for advice on this manuscript, Valley Stewart for advice and guidance, and Samantha Ryken for generating the Δ*tpst* null mutants. This work was funded by the following grants: National Science Foundation (IOS-2436798) to MB and (IOS-1954929) to PCR, National Institute of Health (R35 GM148173) to PCR, USDA-NIFA-AFRI Postdoctoral Fellowship (2023-67012-39889) to AMS, and Dartmouth Undergraduate Research (Barbara E. Crute Memorial Internship) to DVT.

## Supplementary figure legends

**Supplementary Figure 1. CRISPR-Cas9-mediated editing of *TPST*.** A) Diagram of two predicted gene models of the *TPST* locus. Coding exons are represented by green (V3.5) or magenta (V3.1) boxes. Green lines represent the 5’ and 3’ UTR. Location of the protospacer targeted to create a *tpst* null mutant is denoted by PS1. The sequences below show the sequence around PS1, with the PS1 sequence in red. The PAM is underlined. All mutants have the indicated mutations, resulting in early translational stops. Deletions are indicated by dashes, and insertions are indicated in green text. Small arrows above each model indicate primers used for RT-PCR. Numbers on the right of each model indicate product size using cDNA. B) PCR products obtained with the indicated primers using either genomic DNA or cDNA isolated from protonemal tissue were separated on an agarose gel. Green arrow indicates the V3.5 transcript, and magenta arrow indicates the V3.1 transcript. C) TPST predicted primary structure encoded by the V3.5 or V3.1 transcript, drawn approximately to scale. SP denotes the signal peptide sequence. PAPs binding denotes the region between the first residue of the 5’-PSB motif and the last residue of the 3’-PB motif. Black lines indicate the approximate position of the 5’-PSB and 3’-PB elements, and the catalytic base. TM denotes the predicted transmembrane domain. The number of amino acids between the last residue of the 3’-PB and the first residue of the transmembrane domain is indicated below the dashed lines. D) Representative brightfield and chloroplast autofluorescence images of wild type, Δ*tpst-6*, and Δ*tpst-7* from 2-week-old plants that were regenerated from protoplasts on cellophane. Scale bar, 500 µm. E) Plant area was quantified and normalized to wild type. Significant differences determined by a Kruskal-Wallis test (α = 0.05) with a Dunn’s post hoc test and Bonferroni correction are indicated by different letters (n = 20). F and G) Quantification of protonemal growth rate (F) and subapical cell length (G) of caulonemal and chloronemal filaments from 2-week-old plants regenerated from protoplasts. Statistically significant difference in means was calculated using a Student’s t-test for unpaired data with unequal variance (α = 0.05) and is indicated by different letters. H) Quantification of gametophore and bud formation of 2-week-old ground tissue. Significant differences determined by a Kruskal-Wallis test (α = 0.05) with a Dunn’s post hoc test and Bonferroni correction are indicated by different letters.

**Supplementary Figure 2. Δ*tpst* undergoes fewer cell divisions during bud development.** A) Diagram of the two predicted gene models of the *TPST* locus. Coding exons are represented by green (V3.5) or magenta (V3.1) boxes. Green lines represent the 5’ and 3’ UTR. Location of the protospacer targeted to create a *tpst* null mutant is denoted by PS1. The sequences below show the sequence around PS1, with the PS1 sequence in red. The PAM is underlined. Insertions are indicated in green text. B and C) Confocal time-lapse images of wild type (B) and Δ*tpst* (C) of the same buds depicted in Fig. 2B. Each image represents a different cell division at the corresponding time. Arrowheads denote a cell division marked by phragmoplast expansion with mEGFP-tubulin. Scale bar for all images, 20 µm. Also see Videos 1 and 2.

**Supplementary Figure 3. Creating point mutations in *TPST*.** Diagram of the *TPST* locus. Coding exons are represented by green boxes. Black lines represent the coding region, and green boxes represent the 5’ and 3’ UTR. Location of the protospacers used to create point mutations through homology-directed repair is denoted by PS1-3. The sequences below show the protospacer sequence in red, and the PAMs are underlined. Lowercase blue letters indicate base pair changes to incorporate silent mutations or alanine substitutions in bold. Blue capital letters represent the point mutation.

**Supplemental Figure 4: Synthetic PSY1 peptide from three distinct plant species promotes root elongation in rice seedlings.** Quantification of primary root elongation of Kitaake seedlings (n > 20) grown hydroponically in 1X MS with or without indicated synthetic peptide treatment (500nM), 6-days-post treatment. Different letters indicate significant differences determined by a Kruskal-Wallis test (α = 0.05) with a Dunn’s post hoc test.

## Video Legends

**Video 1. Early cell divisions during wild-type bud formation.** Time-lapse images of a wild-type bud growing in a microfluidic device. Plasma membranes are labeled with 3XmRuby2-TON1(magenta), and microtubules are labeled with mEGFP-tubulin (green). Images are maximum projections of z-stacks acquired every 10 minutes on a laser scanning confocal microscope. Video is playing at 10 fps. Scale bar, 20 μm. See also Figure 2B and Supplementary Figure 2B.

**Video 2. Early cell divisions during** Δ***tpst* bud formation.** Time-lapse images of a Δtpst bud growing in a microfluidic device. Plasma membranes are labeled with 3XmRuby2-TON1(magenta), and microtubules are labeled with mEGFP-tubulin (green). Images are maximum projections of z-stacks acquired every 10 minutes on a laser scanning confocal microscope. Video is playing at 10 fps. Scale bar, 20 μm. See also Figure 2B and Supplementary Figure 2C.

**Video 3. Bud development in wild type.** Time-lapse images of the same wild-type bud in Video 1 growing in a microfluidic device. Images are maximum projections of z-stacks acquired every 10 minutes on a laser scanning confocal microscope. Video is playing at 20 fps. Scale bar, 20 μm. See also Figure 2C.

**Video 4. Bud development in** Δ***tpst*.** Time-lapse images of the same Δ*tpst* bud in Video 2 growing in a microfluidic device. Images are maximum projections of z-stacks acquired every 10 minutes on a laser scanning confocal microscope. Video is playing at 20 fps. Scale bar, 20 μm. See also Figure 2C.

**Video 5. Phyllid expansion in a developing wild-type bud.** Time-lapse images of a wild-type bud growing in a microfluidic device. Plasma membranes are labeled with 3XmRuby2-TON1(magenta), and microtubules are labeled with mEGFP-tubulin (green). Images are maximum projections of z-stacks acquired every 10 minutes on a laser scanning confocal microscope. Video is playing at 20 fps. Scale bar, 20 μm. See also Figure 2D.

**Video 6. Phyllid expansion in a developing** Δ***tpst* bud.** Time-lapse images of a Δ*tpst* bud growing in a microfluidic device. Plasma membranes are labeled with 3XmRuby2-TON1(magenta), and microtubules are labeled with mEGFP-tubulin (green). Images are maximum projections of z-stacks acquired every 10 minutes on a laser scanning confocal microscope. Video is playing at 20 fps. Scale bar, 20 μm. See also Figure 2D.

**Video 7. Wild-type gametophore expansion.** Time-lapse images of a wild-type gametophore growing in a microfluidic device. Plasma membranes are labeled with 3XmRuby2-TON1(magenta), and microtubules are labeled with mEGFP-tubulin (green). Images are maximum projections of z-stacks acquired every 10 minutes on a laser scanning confocal microscope. Video is playing at 20 fps. Scale bar, 20 μm. See also Figure 3A.

**Video 8. Gametophore expansion in** Δ***tpst*.** Time-lapse images of a Δ*tpst* gametophore growing in a microfluidic device. Plasma membranes are labeled with 3XmRuby2-TON1(magenta), and microtubules are labeled with mEGFP-tubulin (green). Images are maximum projections of z-stacks acquired every 10 minutes on a laser scanning confocal microscope. Video is playing at 20 fps. Scale bar, 20 μm. See also Figure 3A.

## Supplementary tables

**Table S1.** Mutant lines used and created.

| Name | Background | Genetic alteration | Translation |
| --- | --- | --- | --- |
| $\Delta tpst-6$ ($\Delta tpst$) | Wild type | 7 bp deletion | Stop at AA90, aa 81-89 are irrelevant |
| $\Delta tpst-7$ | Wild type | Stop oligo insertion | Stop at AA85, aa 82-84 are irrelevant |
| *tpst-R81A* | Wild type | CGA→ GCT (241-243, AA change), C246T (silent), C249A (silent), C252A (silent) | R81A |
| *tpst-H124A* | Wild type | C348T (silent), C351T (silent), C354T (silent), T361C (silent), G363T (silent), T366A (silent), A369T (silent), CAC→ GCT (370-372, AA change) | H124A |
| *tpst-R144A* | Wild type | CGC→ GCT (430-432, AA change), C435T (silent), A438T (silent), G441T (silent), T444C (silent), T447A (silent) | R144A |
| 3XmRuby2-TON1 | mEGFP-tubulin | Insertion of sequences encoding 3 tandem mRuby2 proteins at the 5' end of the *TON1* gene at the endogenous locus | N-terminal fusion of 3XmRuby2 with TON1 |
| 3XmRuby2-TON1/$\Delta tpst$ | 3XmRuby2-TON1 | Stop oligo insertion | Stop at AA85, aa 82-84 are irrelevant |

**Table S2.**
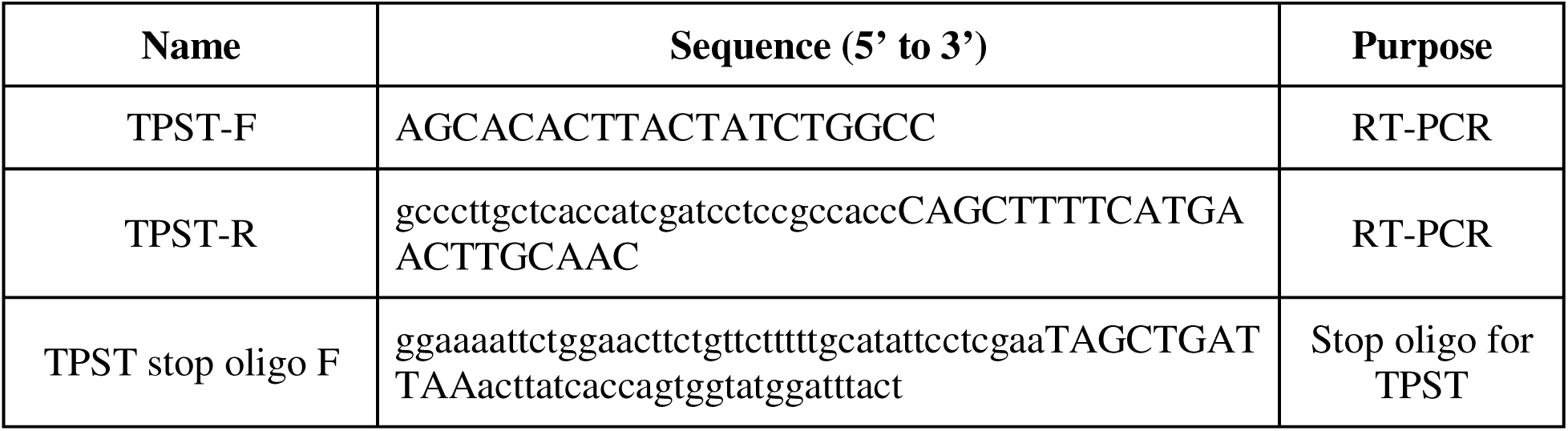
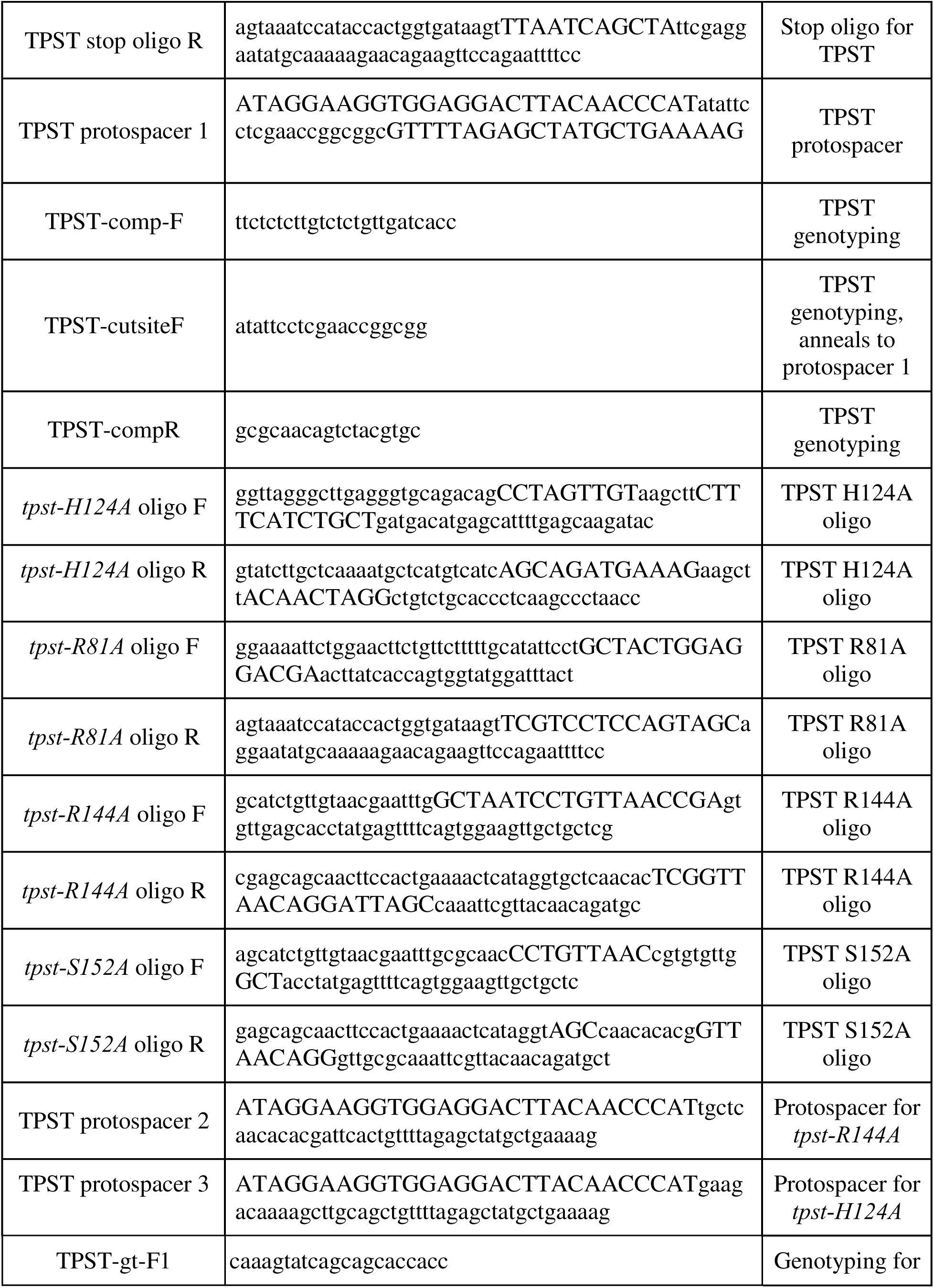
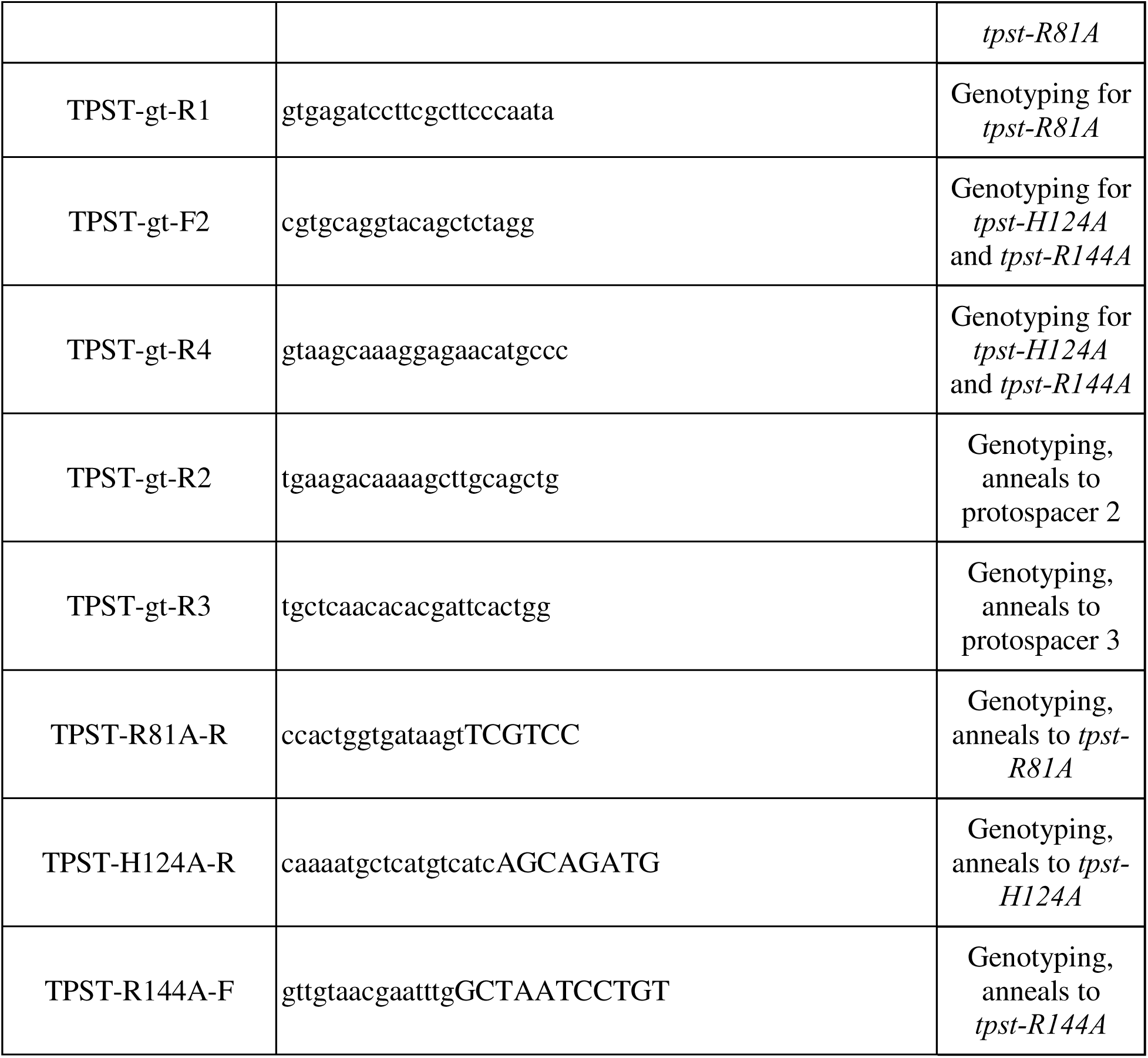
Table of primers, oligos, and protospacer sequences.

