## Supplemental Figure 1 for "Evidence for Early Evolution of Sulfated Peptide Signaling in Plant Development"

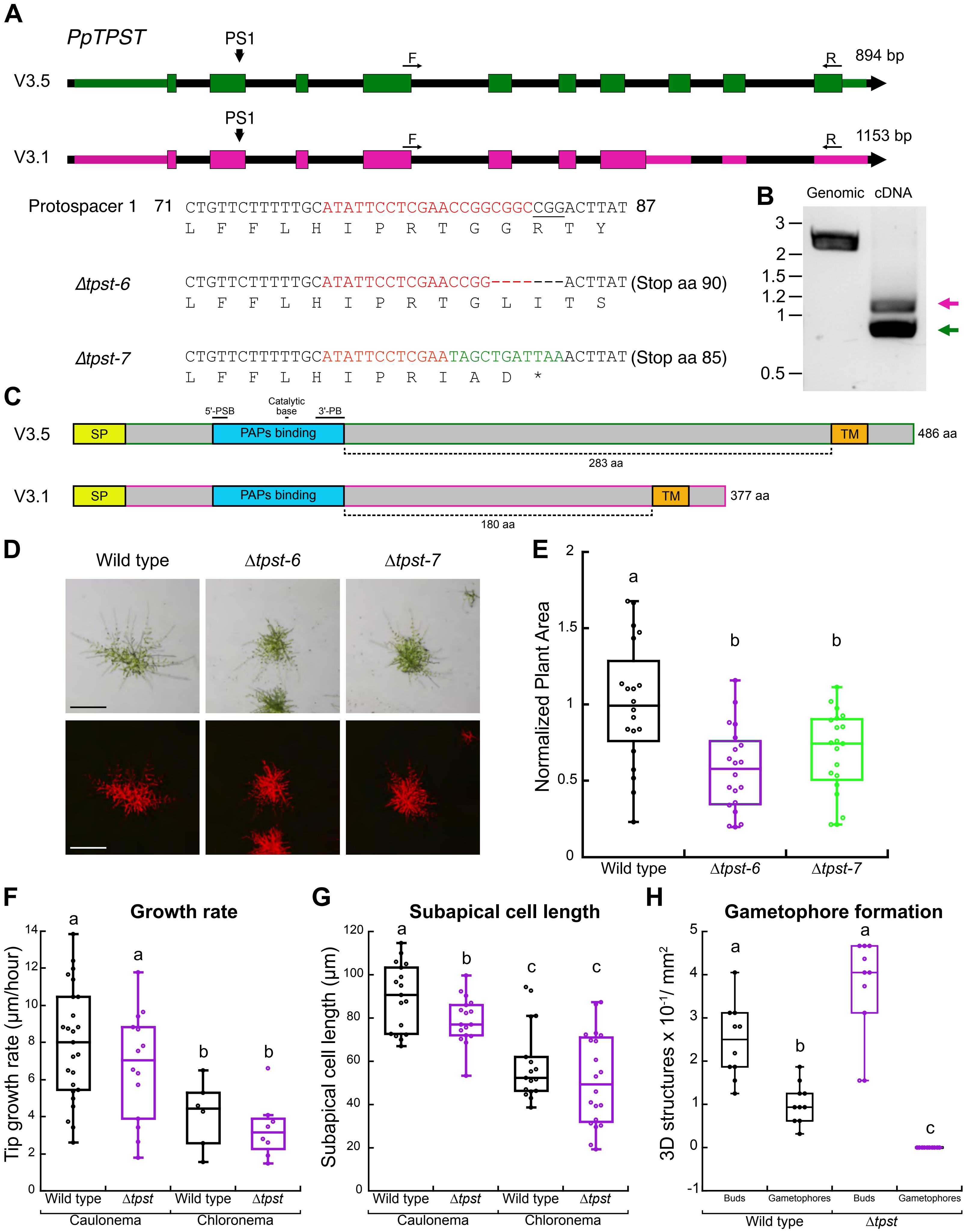

A
PpTPST
PS1
F
R
894 bp
V3.5
V3.1
1153 bp
Protospacer 1 71 CTGTTCTTTTTGCATATTCCTCGAACCGGCGGCCGGACTTAT 87
L F F L H I P R T G G R T Y
Δtpst-6 CTGTTCTTTTTGCATATTCCTCGAACCGG-------ACTTAT (Stop aa 90)
L F F L H I P R T G L I T S
Δtpst-7 CTGTTCTTTTTGCATATTCCTCGAATAGCTGATTAAACTTAT (Stop aa 85)
L F F L H I P R I A D *
B
Genomic cDNA
3
2
1.5
1.2
1
0.5
C
5'-PSB
Catalytic base
3'-PB
V3.5
SP
PAPs binding
TM
486 aa
283 aa
V3.1
377 aa
180 aa
D
Wild type
Δtpst-6
Δtpst-7
E
Normalized Plant Area
a
b
b
Wild type
Δtpst-6
Δtpst-7
F
Growth rate
Tip growth rate (μm/hour)
Wild type
Δtpst
Caulonema
Chloronema
G
Subapical cell length
Subapical cell length (μm)
H
Gametophore formation
3D structures x $10^{-1}$/ $mm^2$
Buds
Gametophores
