## Supplemental Figure 2 for "Evidence for Early Evolution of Sulfated Peptide Signaling in Plant Development"

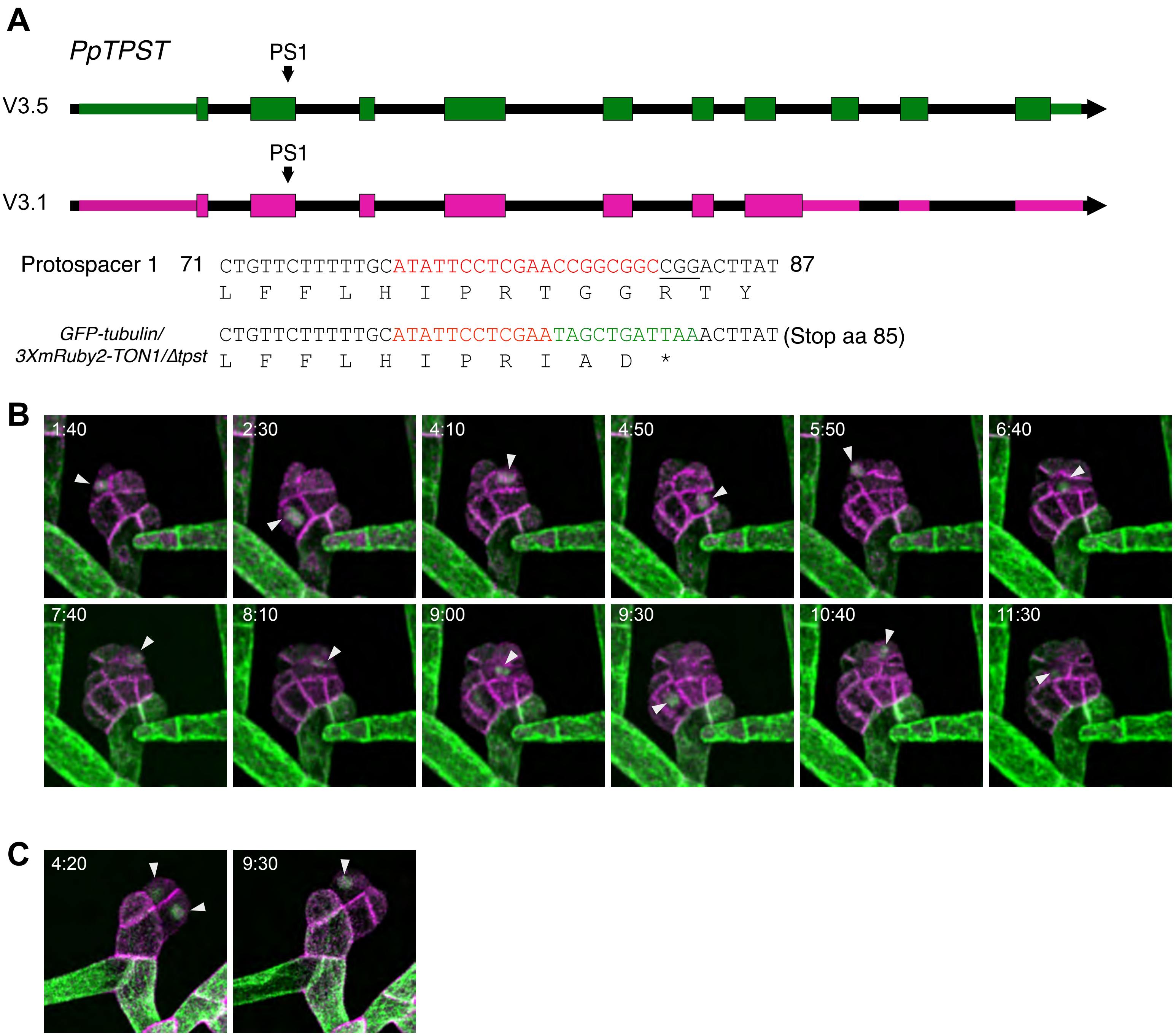


**Supplementary Figure 2. Δ*tpst* undergoes fewer cell divisions during bud development.** A) Diagram of the two predicted gene models of the *TPST* locus. Coding exons are represented by green (V3.5) or magenta (V3.1) boxes. Green lines represent the 5' and 3' UTR. Location of the protospacer targeted to create a *tpst* null mutant is denoted by PS1. The sequences below show the sequence around PS1, with the PS1 sequence in red. The PAM is underlined. Insertions are indicated in green text. B and C) Confocal time-lapse images of wild type (B) and Δ*tpst* (C) of the same buds depicted in Fig. 2B. Each image represents a different cell division at the corresponding time. Arrowheads denote a cell division marked by phragmoplast expansion with mEGFP-tubulin. Scale bar for all images, 20 μm. Also see Videos 1 and 2.
