## Supplemental Figure 3 for "Evidence for Early Evolution of Sulfated Peptide Signaling in Plant Development"

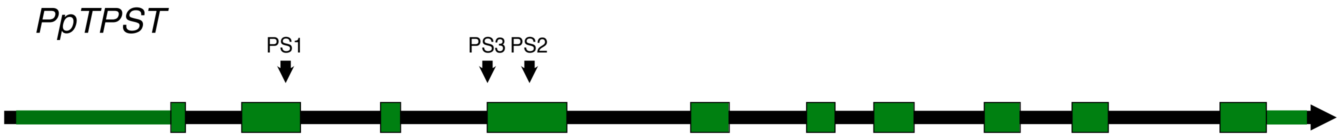


Protospacer 1 74 CTGTTCTTTTTGCATATTCCTCGAACCGGCGGCCGGACTTAT 87
L F F L H I P R T G G R T Y

*tpst*-R81A CTGTTCTTTTTGCATATTCCT**gct**ACtGGaGGaCGaACTTAT
L F F L H I P **A** T G G R T Y

Protospacer 2 142 AATTTGCGCAACCCAGTGAATCGTGTGTTGAGCACCTATGAG 155
N L R N P V N R V L S T Y E

*tpst*-R144A AATTTG**gct**AAtCCtGTtAAcCGaGTGTTGAGCACCTATGAG
N L **A** N P V N R V L S T Y E

Protospacer 3 115 CAGCCCAGCTGCAAGCTTTTGTCTTCACACGATGACATGAGC 128
Q P S C K L L S S H D D M S

*tpst*-H124A CAGCCtAGtTGtAAGCTTcTtTCaTCt**gct**GATGACATGAGC
Q P S C K L L S S **A** D D M S

**Supplementary Figure 3. Creating point mutations in *TPST*.** Diagram of the *TPST* locus. Coding exons are represented by green boxes. Black lines represent the coding region, and green boxes represent the 5' and 3' UTR. Location of the protospacers used to create point mutations through homology-directed repair is denoted by PS1-3. The sequences below show the protospacer sequence in red, and the PAMs are underlined. Lowercase blue letters indicate base pair changes to incorporate silent mutations or alanine substitutions in bold. Blue capital letters represent the point mutation.
