## Supplemental Figure 4 for "Evidence for Early Evolution of Sulfated Peptide Signaling in Plant Development"

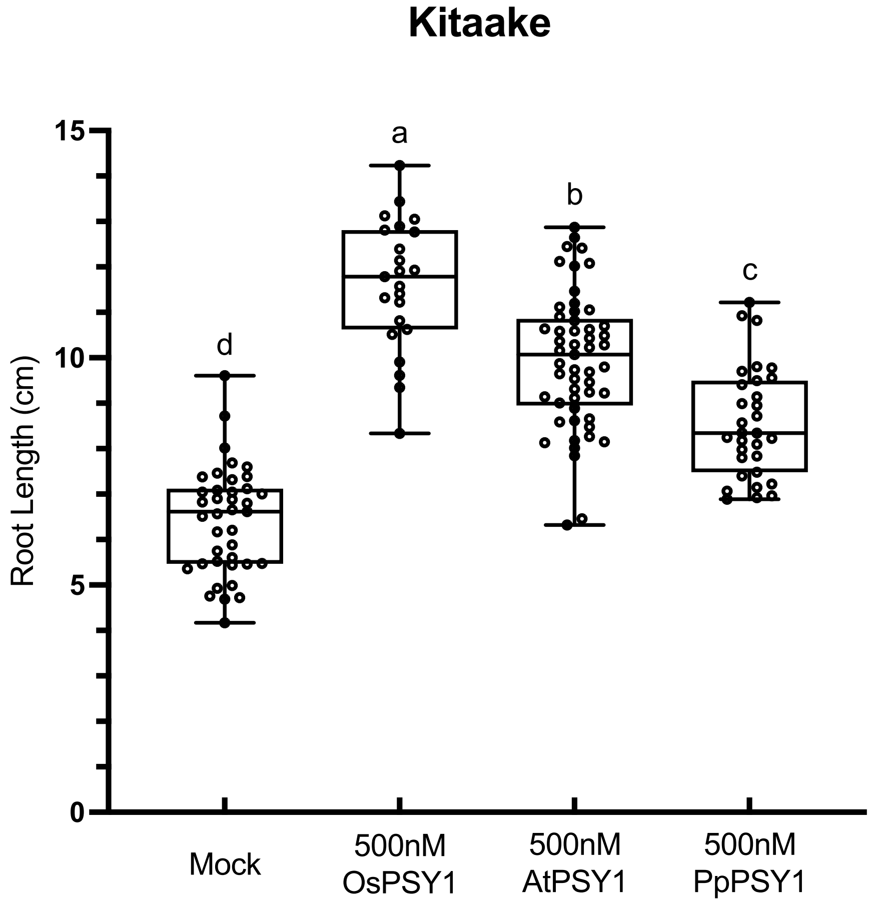


**Supplemental Figure 4: Synthetic PSY1 peptide from three distinct plant species promotes root elongation in rice seedlings.** Quantification of primary root elongation of Kitaake seedlings (n  20) grown hydroponically in 1X MS with or without indicated synthetic peptide treatment (500nM), 6-days-post treatment. Different letters indicate significant differences determined by a Kruskal-Wallis test (α = 0.05) with a Dunn's post hoc test.
